## Supplementary Information for "Intrinsic timescales in the visual cortex change with selective attention and reflect spatial connectivity"

**2 Supplementary tables** **13**

**3 Supplementary figures** **15**

### 1 Supplementary notes

#### 1.1 Relation between modulation of timescales and power spectra during attention

Attention is known to affect the spectral power of neural activity fluctuations at different frequencies. Previous studies reported a reduction in the low-frequency power ($\sim 2-20$ Hz) of spike-triggered average local field potentials (LFPs) [1], spike-LFP coherence [2], and spike-spike coherence[3] during attention in area V4. In addition, an increase in the gamma frequency band ($\sim 40-60$ Hz) was detected in spike-triggered average LFPs [1], spike-LFP coherence [2], and LFP spectral coherence [4]. Similar changes in the low-frequency and gamma-range LFP power were also observed in our data during attention task 1 [5]. We also observed a reduction at low-frequencies in the power spectral density (PSD) of spiking activity in our data (Supplementary Fig. 6) as in previous studies [3].

To test how these changes in the PSD shape are related to modulation of timescales during attention, we estimated timescales directly from PSD using our aABC method in the frequency domain [6]. We fitted the PSD of spiking activity with the two-timescale generative model and obtained the same results as with the time-domain analyses: the fast timescale did not change and the slow timescale increased with attention (Supplementary Fig. 7, cf. Fig. 3b, e). These results show that the increase of the slow timescale is consistent with the reduction in the power of low-frequency fluctuations during attention.

To explain the co-occurrence of these two phenomena, we considered theoretically how the PSD shape changes under the modulation of timescales. In the frequency domain, PSD of a one-timescale process with an exponential autocorrelation is a Lorentzian function $\frac{c}{f^2+f_k^2}$, where $f$ is the frequency, $f_k$ is the knee frequency related to the timescale $\tau$ as $f_k = (2\pi\tau)^{-1}$, and $c$ is a normalization constant. In logarithmic-logarithmic coordinates this function has a plateau at low frequencies transitioning to the $1/f^2$ decay at the knee frequency. PSD of a two-timescale process is a mixture of two Lorentzian functions, one with a large knee frequency (fast timescale) and one with a small knee frequency (slow timescale). In V4 data, the slow timescale increased and the fast timescale did not change during attention. When the slow timescale increases, the small knee frequency shifts to even lower frequencies. At the same time, the large knee frequency (the fast timescale) does not change, keeping the power at high frequencies constant. For the values of timescales estimated in V4, these effects together produce a reduction in the low-frequency power ($\sim 2-20$ Hz, Supplementary Fig. 6). The decrease in the small knee frequency indeed results in an increase of power at very low frequencies ($< 2$ Hz), which, however, is not detectable with short observation windows ($< 500$ ms) used in previous studies. With a longer observation window of 700 ms, we indeed see a PSD increase at a very low frequency (1.4 Hz) during attention (Supplementary Fig. 6).

#### 1.2 Number of timescales in single unit activity

We tested whether multiple timescales can also be observed in autocorrelations of the single-unit activity (SUA). However, due to low firing rates, SUA often does not yield sufficient data for conclusive model comparison, therefore, we used multi-unit activity (MUA) for the rest of analyses. We fitted autocorrelations of SUA during the fixation task (which had the longest trial duration of 3 s and thus the largest data amount) and performed the model comparison to determine the number of timescales (Supplementary Fig. 8). For these analyses the autocorrelations were fitted from the time-lag $t > 0$ with the largest correlation value to discard the effects related to the refractory period or the negative adaptation in initial time lags (similar to [7]).

While some single units clearly showed two distinct timescales, the model comparison was inconclusive for most units because autocorrelations were dominated by noise due to low data amount. In addition, some SUA autocorrelations exhibited more complex dynamics (e.g., oscillations or refractory period) than what can be captured by the one- or two-timescale model. Due to these limitations of SUA (also reported by previous studies [8, 9]), it is not possible to conclusively perform the analyses of timescales and their attentional modulation using SUA.

#### 1.3 Autocorrelation of a two-state Markov process

We consider a Markov process (2SMP) with two states $\{0, 1\}$ and the transition matrix

$$\mathbb{P} = \begin{bmatrix} \omega & 1-\omega \\ 1-\theta & \theta. \end{bmatrix} \tag{1}$$

We can compute the autocorrelation of the discrete-time time-series $X_n$ in this model as

$$\begin{aligned} AC(t) &= \mathbb{E}[\,X_n X_{n+t}] - \mathbb{E}[\,X_n]\,\mathbb{E}[\,X_{n+t}] && (2)\\ &= \mathbb{E}[\,X_n X_{n+t}|X_n = 1]\,P(X_n = 1) && (3)\\ &\quad + \mathbb{E}[\,X_n X_{n+t}|X_n = 0]\,P(X_n = 0) && (4)\\ &\quad - P(X_n = 1)P(X_{n+t} = 1) && (5)\\ &= P(X_n = 1)\left[P(X_{n+t} = 1|X_n = 1) - P(X_{n+t} = 1)\right]. && (6) \end{aligned}$$

Here $n$ is the time-step, $t$ is the time-lag and $\mathbb{E}[.]$ gives the expectation value.

For the Markov process we can write

$$P(X_n = 1) = P(X_{n+t} = 1) = \pi_1, \tag{7}$$

where $\pi_1$ is the stationary distribution of state 1. The stationary distribution $\pi$ of 2SMP is invariant with respect to the transition matrix

$$\pi\mathbb{P} = \pi, \tag{8}$$

where $\pi$ is

$$\pi = \begin{bmatrix} \pi_0 & \pi_1. \end{bmatrix} \tag{9}$$

By transposing Eq.(8) we get

$$\mathbb{P}^{\intercal}\pi^{\intercal} = \pi^{\intercal}. \tag{10}$$

This equation shows that the stationary distribution $\pi$ is the eigenvector of $\mathbb{P}^{\intercal}$ for eigenvalue of 1.

The eigenvalue equation can be written as

$$\begin{bmatrix} \omega & 1-\theta \\ 1-\omega & \theta \end{bmatrix} \begin{bmatrix} \pi_0 \\ \pi_1 \end{bmatrix} = \begin{bmatrix} \pi_0 \\ \pi_1 \end{bmatrix}. \tag{11}$$

In addition, $\pi$ satisfies the normalization condition for a probability distribution

$$\pi_0 + \pi_1 = 1. \tag{12}$$

From Eq.(11) and Eq.(12) we compute the stationary distributions as

$$\pi_0 = \frac{1-\theta}{2-\omega-\theta}, \tag{13}$$

$$\pi_1 = \frac{1-\omega}{2-\omega-\theta}. \tag{14}$$

To find the autocorrelation, we also need to compute the transition probability $P(X_{n+t} = 1|X_n = 1)$. This term is the probability of observing state 1, when we observed state 1 at $t$ time-steps before. This transition probability for $t$ time-steps is determined by $\mathbb{P}^t$.

To calculate $\mathbb{P}^t$, we use its eigenvalue decomposition

$$\mathbb{P}^t = V\Lambda^t V^{-1}, \tag{15}$$

where

$$\Lambda^t = \begin{bmatrix} 1 & 0 \\ 0 & (\omega+\theta-1)^t \end{bmatrix} \tag{16}$$

and

$$V = \begin{bmatrix} \frac{1}{\sqrt{2}} & \frac{1}{\sqrt{1+\frac{1-\omega}{1-\theta}^2}} \\ \frac{1}{\sqrt{2}} & \frac{1-\omega}{\sqrt{(1-\omega)^2+(1-\theta)^2}}. \end{bmatrix} \tag{17}$$

From these equations we can compute

$$P(X_{n+t} = 1|X_n = 1) = \mathbb{P}^t_{2,2} \tag{18}$$

Finally, using Eq.(6), Eq.(14) and Eq.(18) we calculate the autocorrelation function of 2SMP

$$\mathcal{AC}(t) = \frac{(1-\omega)(1-\theta)}{(2-\omega-\theta)}(\omega+\theta-1)^t. \tag{19}$$

#### 1.4 Analytical derivations of the timescales in the spatial network model

Here we analytically derive the timescales for the dynamics in the spatial network model (details in [10]). The network model is defined on a square lattice with $N' \times N'$ nodes with periodic boundary conditions. The side length of the lattice is $L$, and the distance between two neighboring nodes is $L/N' = a$ (considering the boundary conditions). Each unit is located at one node, with two-dimensional coordinates $\mathbf{x} = (x_1, x_2)$, where $x_1 = m_1 L/N'$, $x_2 = m_2 L/N'$, $(m_{1,2} = 0, \ldots, N-1')$. In the locally connected network, each unit receives inputs from its nearest neighbors ($n = 8$).

Activity of each unit $i$ is described by a binary state variable $S_i \in \{0, 1\}$. The state of each unit $S_i$ at time-step $t'$ is updated according to the transition rates

$$\begin{aligned} w(S_i = 0 \to 1) &= \alpha_1 + \beta_1\, \mathcal{F}\left(\sum_j S_j\right), \\ w(S_i = 1 \to 0) &= \alpha_2 - \beta_2\, \mathcal{F}\left(\sum_j S_j\right). \end{aligned} \tag{20}$$

Here $\mathcal{F}(\sum_j S_j)$ defines the interaction term between the neighbors. For the linear model, the interaction term is proportional to the number of active neighbors at time $t'$, and we can relate the transition rates to the parameters of the discrete-time version of the model: $\alpha_1 = p_{\text{ext}} \left[\frac{-\ln(p_s)}{(1-p_s)\Delta t'}\right]$ and $\alpha_2 = (1 - p_s - p_{\text{ext}}) \left[\frac{-\ln(p_s)}{(1-p_s)\Delta t'}\right]$, $\beta_1 = \beta_2 = p_r \left[\frac{-\ln(p_s)}{(1-p_s)\Delta t'}\right]$, where $\Delta t'$ is the duration of each discrete time step. In the following, we derive the timescales for a general form of dynamics defined in Eq.(20).

At time $t'$, the probability of the units to be in a specific configuration $\{S\} = \{S_1, S_2, ..., S_N\}$ is denoted by $P(\{S\}, t')$. The master equation describes the time evolution of $P(\{S\}, t')$, that is given by [11]

$$\frac{d}{dt'}P(\{S\},t') = -P(\{S\},t')\sum_i w(S_i) + \sum_i P(\{S\}^{i*},t')w(1-S_i), \tag{21}$$

where $\{S\}^{i*} = \{S_1, S_2, ..., 1 - S_i, ..., S_N\}$.

Using the master equation, we can derive the equations for the time evolution of different moments. For example, the mean activity of the unit $S_i$ is defined as

$$\langle S_i\rangle(t') = \sum_{\{S\}} P(\{S\},t')S_i \,, \tag{22}$$

where we are summing over all configurations of variables $\{S\}$ at a given time $t'$. The time evolution of the mean activity is given by

$$\frac{d}{dt'}\langle S_i\rangle(t') \;=\; \frac{d}{dt'}\left(\sum_{\{S\}} P(\{S\},t')S_i\right) = \sum_{\{S\}}\left(\frac{d}{dt'}P(\{S\},t')\right)S_i \,. \tag{23}$$

Substituting the master equation, we have

$$\frac{d}{dt'}\langle S_i\rangle(t') = \sum_{\{S\}} P(\{S\},t')[w(S_i)(1-2S_i)] \,. \tag{24}$$

According to Eq.(22) and Eq.(20), the time-evolution equation of the mean activity is

$$\frac{d}{dt'}\langle S(\mathbf{x})\rangle = \alpha_1 - (\alpha_1+\alpha_2)\langle S(\mathbf{x})\rangle + \beta_1\left\langle \sum_{\mathbf{x}\pm} S(\mathbf{x})\right\rangle + (\beta_2-\beta_1)\left\langle S(\mathbf{x})\sum_{\mathbf{x}\pm} S(\mathbf{x})\right\rangle \,. \tag{25}$$

Here and below, $\sum_{\mathbf{x}\pm}$ denotes the summation over units connected to the target unit at $\mathbf{x}$. By setting $\frac{d}{dt'}\langle S(\mathbf{x})\rangle = 0$ and averaging over all units, we obtain the steady-state solution for the mean activity $\bar{S}$. In the case of $\beta_1 = \beta_2$, the steady-state mean activity is

$$\bar{S} = \frac{1}{N}\sum_i \langle S_i\rangle = \frac{\alpha_1}{\alpha_1+\alpha_2-n\beta_1} \,. \tag{26}$$

Similarly, the rate of change of average activity of a pair of units is

$$\frac{d}{dt'}\langle S_iS_j\rangle(t') = \sum_{\{S\}} P(\{S\},t')w(S_i)(1-2S_i)S_j + w(S_j)(1-2S_j)S_i. \tag{27}$$

The time evolution of time-delayed quadratic moment at time-lag $t$ is defined as

$$\frac{d}{dt}\langle S_i(t')S_j(t'+t)\rangle = \sum_{\{S\}} P(\{S\},t')S_i \frac{d}{dt}\left(\sum_{\{\sigma\}} P(\{\sigma\},t'+t|\{S\},t')\sigma_j\right) . \tag{28}$$

Here $P(\{\sigma\},t'+t|\{S\},t')$ is the conditional probability of finding the system in configuration $\{\sigma\}$ at time $t'+t$, given that it was in configuration $\{S\}$ at time $t'$. Since the conditional probability obeys the same master equation, we have

$$\frac{d}{dt}\langle S_i(t')S_j(t'+t)\rangle = \langle S_i(t')(1-2S_j(t'+t))w(S_j(t'+t))\rangle . \tag{29}$$

We define $CC(\mathbf{\Delta},t)$ to be the average cross-correlation between two units at coordinates $\mathbf{x}$ and $\mathbf{y}$ with the fixed distance $\mathbf{\Delta} = (\Delta_1, \Delta_2)$, $\Delta_1 = |x_1 - y_1|$, $\Delta_1 = |x_2 - y_2|$ :

$$CC(\mathbf{\Delta},t) = \frac{1}{N'^2} \sum_{x_1=0}^{(N'-1)a} \sum_{y_1=0}^{(N'-1)a} \langle \delta S(\mathbf{x},t')\delta S(\mathbf{y},t'+t)\rangle , \tag{30}$$

where $\delta S = S - \bar{S}$. Using Fourier transformation,

$$\delta S(\mathbf{x},t') = \sum_{k_1,k_2=0}^{2\pi(N'-1)/L} \delta S(\mathbf{k},t')\exp(i\mathbf{k}\cdot\mathbf{x}) , \tag{31}$$

$$\delta S(\mathbf{y},t'+t) = \sum_{k'_1,k'_2=0}^{2\pi(N'-1)/L} \delta S(\mathbf{k}',t'+t)\exp(i\mathbf{k}'\cdot\mathbf{y}) , \tag{32}$$

$CC(\mathbf{\Delta},t)$ can be written as

$$CC(\mathbf{\Delta},t) = \frac{1}{N'} \sum_{k_1,k_2=0}^{2\pi(N'-1)/L} \langle \delta S(\mathbf{k},t')\delta S(-\mathbf{k},t'+t)\rangle \times \exp(i\mathbf{k}\cdot\mathbf{\Delta}) . \tag{33}$$

Therefore, we can define the Fourier transformation of the correlation function $\tilde{CC}(k_1,k_2,t)$ to be $\langle \delta S(\mathbf{k},t')\delta S(-\mathbf{k},t'+t)\rangle/N'$. This transformation can be obtained as:

$$\tilde{CC}(k_1,k_2,t) = \frac{4}{N'^2} \sum_{\Delta_1,\Delta_2=0}^{(N'/2-1)a} CC(\mathbf{\Delta},t)e^{-ik_1\Delta_1 - ik_2\Delta_2} . \tag{34}$$

The equal-time cross-correlation defines the initial condition of time-delayed cross-correlation:

$$CC(\mathbf{\Delta}) \equiv CC(\mathbf{\Delta},t=0) , \quad \tilde{CC}(k_1,k_2) \equiv \tilde{CC}(k_1,k_2,t=0) . \tag{35}$$

The Fourier transformation of equal-time cross-correlation is

$$CC(\mathbf{\Delta}) = \sum_{k_1,k_2=0}^{\frac{2\pi(N'/2-1)}{L}} \cos(k_1\Delta_1)\cos(k_2\Delta_2)\tilde{CC}(k_1,k_2) . \tag{36}$$

Similarly, we define the average autocorrelation to be

$$\mathcal{AC}(t) = \frac{1}{N'^2} \sum_{x_1,x_2=0}^{(N'-1)a} \langle \delta S(\mathbf{x},t)\delta S(\mathbf{x},t'+t)\rangle \ . \tag{37}$$

In the networks of binary units, the initial condition of average autocorrelation, $\mathcal{A}(0) = \mathcal{A}(t=0)$, is fixed by the mean activity $\bar{S}$:

$$\mathcal{A}(0) = (1-\bar{S})\bar{S} \ . \tag{38}$$

The time-evolution of equal-time correlation is

$$\begin{aligned}\frac{d}{dt'}\langle \delta S(\mathbf{x},t')\delta S(\mathbf{y},t')\rangle &= -2(\alpha_1+\alpha_2)\langle \delta S(\mathbf{x},t')\delta S(\mathbf{y},t')\rangle \\ &+\beta_1(\langle \sum_{\mathbf{x}\pm} \delta S(\mathbf{x})\delta S(\mathbf{y})\rangle + \langle \delta S(\mathbf{x}) \sum_{\mathbf{y}\pm} \delta S(\mathbf{y})\rangle) \ . \end{aligned} \tag{39}$$

For the steady state, by averaging across $\mathbf{x}$ and fixing distance $\mathbf{\Delta}$, we obtain the equation for the average correlation $CC(\mathbf{\Delta},0)$:

$$\begin{aligned} CC(\mathbf{\Delta}) = \frac{\beta_1}{\alpha_1+\alpha_2}[&CC(\Delta_1-a,\Delta_2)+CC(\Delta_1+a,\Delta_2)+CC(\Delta_1,\Delta_2+a)+CC(\Delta_1,\Delta_2-a) \\ &+CC(\Delta_1+a,\Delta_2+a)+CC(\Delta_1+a,\Delta_2-a)+CC(\Delta_1-a,\Delta_2+a)+CC(\Delta_1-a,\Delta_2-a) \\ &+(\delta_{\Delta_1,0}\delta_{\Delta_2,a}+\delta_{\Delta_1,0}\delta_{\Delta_2,-a}+\delta_{\Delta_1,-a}\delta_{\Delta_2,0}+\delta_{\Delta_1,a}\delta_{\Delta_2,0}+\delta_{\Delta_1,a}\delta_{\Delta_2,a}+\delta_{\Delta_1,a}\delta_{\Delta_2,-a} \\ &+\delta_{\Delta_1,-a}\delta_{\Delta_2,a}+\delta_{\Delta_1,-a}\delta_{\Delta_2,-a})\mathcal{AC}(0)] \ . \end{aligned} \tag{40}$$

Performing Fourier transformation, we obtain the solution of equations for each $\mathbf{k}$ mode:

$$\tilde{C}C(k_1,k_2) = \frac{\frac{2\beta_1}{\alpha_1+\alpha_2}[2\cos(k_1a)\cos(k_2a)+\cos(k_1a)+\cos(k_2a)]}{1-\frac{2\beta_1}{\alpha_1+\alpha_2}[(2\cos(k_1a)\cos(k_2a)+\cos(k_1a)+\cos(k_2a)]} \times \frac{4}{N'^2}\mathcal{AC}(0) \ . \tag{41}$$

W can solve for the correlation function $CC(\mathbf{\Delta})$ by an inverse Fourier transformation of $\tilde{C}_2(k_1,k_2)$:

$$CC(\mathbf{\Delta}) = \mathcal{AC}(0)\exp\left(-\frac{\Delta_1+\Delta_2}{\xi}\right) \ . \tag{42}$$

Here $\xi$ is defined as the correlation length:

$$\xi = a\cdot\frac{1}{\ln\left(f'+\sqrt{f'^2-1}\right)} \ , \qquad f' = -\frac{1}{2}+\frac{1}{2}\sqrt{1+\frac{\alpha_1+\alpha_2}{\beta_1}} \ . \tag{43}$$

By using the $CC(\mathbf{\Delta})$ as an initial condition, we can slove for the time-delayed cross-correlation $CC(\mathbf{\Delta},t)$. The evolution equation for the time-delayed cross-correlation is

$$\begin{aligned} &\tau_{\text{self}}\frac{d}{dt}\langle \delta S(\mathbf{x},t')\delta S(\mathbf{y},t'+t)\rangle = \\ -\ &\langle \delta S(\mathbf{x},t')\delta S(\mathbf{y},t'+t)\rangle + \frac{\beta_1}{\alpha_1+\alpha_2}(\langle \delta S(\mathbf{x},\mathbf{t}')\sum_{\mathbf{y}\pm}\delta S(\mathbf{y},\mathbf{t}'+\mathbf{t})\rangle) \ , \end{aligned} \tag{44}$$

where $\tau_{\text{self}}$ is the self-excitation timescale:

$$\tau_{\text{self}} = \frac{1}{\alpha_1 + \alpha_2} \,. \tag{45}$$

Averaging across all $\mathbf{x}$ and fixing distance $\mathbf{\Delta}$, we obtain the equation for the average cross-correlation $CC(\mathbf{\Delta}, t)$. We then apply Fourier transform to the above equation. To further simplify the equation, we neglect the contribution of $\mathcal{AC}(t)$ at the right-hand side of the equation. With this approximation, we find the time evolution of each Fourier mode $\tilde{C}C(k_1, k_2, t)$ that is described by an interaction timescale:

$$\tilde{C}C(k_1, k_2, t) = \tilde{C}C(k_1, k_2) \exp\left(-\frac{t}{\tau_{\text{int},\mathbf{k}}\,(k_1, k_2)}\right) \,, \tag{46}$$

where the interaction timescale is

$$\tau_{\text{int},\mathbf{k}}\,(k_1, k_2) = \frac{\tau_{\text{self}}}{1 - \frac{2\beta_1}{\alpha_1+\alpha_2}[2\cos(k_1 a)\cos(k_2 a) + \cos(k_1 a) + \cos(k_2 a)]} \,. \tag{47}$$

The largest interaction timescale is given by $\mathbf{k} = 0$:

$$\tau_{\text{global}} = \tau_{\text{int},\mathbf{k}}\,(k_1 = 0, k_2 = 0) = \frac{\tau_{\text{self}}}{1 - \frac{n\beta_1}{\alpha_1+\alpha_2}} = \frac{1}{\alpha_1 + \alpha_2 - n\beta_1} \,. \tag{48}$$

The time evolution of average cross-correlation $CC(\mathbf{\Delta}, t)$ is given by a superposition of $N'^2/4$ modes, where each mode has an interaction timescale $\tau_{\text{int},\mathbf{k}}\,(k_1, k_2)$:

$$CC(\mathbf{\Delta}, t) = \sum_{k_1, k_2 = 0}^{\frac{2\pi(N'/2-1)}{L}} \tilde{C}C(k_1, k_2)\cos(k_1\Delta)\cos(k_2\Delta)\exp\left(-\frac{t}{\tau_{\text{int},\mathbf{k}}\,(k_1, k_2)}\right) \,. \tag{49}$$

The time-evolution of autocorrelation is governed by

$$\begin{aligned} &\tau_{\text{self}}\frac{d}{dt}\langle \delta S(\mathbf{x}, t')\delta S(\mathbf{x}, t'+t)\rangle = \\ -\;\; &\langle \delta S(\mathbf{x}, t')\delta S(\mathbf{x}, t'+t)\rangle + \frac{\beta_1}{\alpha_1+\alpha_2}(\langle \delta S(\mathbf{x}, \mathbf{t}')\sum_{\mathbf{x}\pm}\delta S(\mathbf{x}, \mathbf{t}'+\mathbf{t})\rangle) \,. \end{aligned} \tag{50}$$

Averaging over $\mathbf{x}$, we obtain

$$\tau_{\text{self}}\frac{d}{dt}\mathcal{AC}(t) = -\mathcal{AC}(t) + \frac{\beta_1}{\alpha_1+\alpha_2}\left[\sum_{\mathbf{m}} CC(\mathbf{m}, t)\right] \,. \tag{51}$$

where $\mathbf{m} = (a, a),\ (-a, a),\ (a, -a),\ (-a, -a),\ (a, 0),\ (-a, 0),\ (0, a),\ (0, -a)$. The exact solution of this equation is given by

$$\mathcal{AC}(t) = \mathcal{AC}(0)\exp(-\frac{t}{\tau_{\text{self}}}) + \beta_1\exp(-\frac{t}{\tau_{\text{self}}})\int_0^t \sum_{\mathbf{m}} CC(m, t)\exp(\frac{t'}{\tau_{\text{self}}})dt' \,, \tag{52}$$

Substituting the analytical solution of $CC(\Delta, t)$, we have the analytical form of autocorrelation:

$$\mathcal{AC}(t) = \mathcal{AC}(0)\exp\left(-\frac{t}{\tau_{\text{self}}}\right) + \sum_{k_1, k_2 = 0}^{\frac{2\pi(N'/2-1)}{L}} \tilde{C}C(k_1, k_2)\left[\exp\left(-\frac{t}{\tau_{\text{int},\mathbf{k}}\,(k_1, k_2))}\right)\right] \,. \tag{53}$$

#### 1.5 Local timescales are shaped by the spatial network structure

Since the model with spatial connectivity could best explain the V4 spatiotemporal correlations, we further investigated how the timescales of local dynamics depend on the structure of spatial connectivity. We systematically modified the range of recurrent connections between units in the model, while keeping the strength and number of connections constant. We fixed a connectivity radius $r$ and connected each unit to its $8$ randomly chosen neighbors within this radius (Fig. 6a). For small $r$ in the model, the connections between units are local (as in Fig. 4c). With increasing $r$, the connections become more dispersed, and when $r$ reaches the network size, we get a randomly connected network.

We found that timescales of local dynamics reflect the underlying spatial network structure (Fig. 6b). Autocorrelations ($AC$) measured from the activity of individual units decay faster in networks with more dispersed connectivity compared to locally connected networks. In addition, the weight of interaction timescales in the autocorrelation decreases with increasing connectivity radius. As a result, the transition point between the self-excitation and interaction timescales shifts towards longer time-lags so that the self-excitation timescale dominates the autocorrelation in networks with dispersed connectivity.

While networks with local versus dispersed connectivity generate distinct local timescales, the timescale of their global dynamics is similar (Fig. 6b, inset). The timescale of global network activity is a direct measure for distance of system's dynamics from the critical point, with longer timescales observed in systems closer to criticality [12]. We measured the global timescale from the autocorrelation of the activity summed across all units in the network ($AC_{\text{global}}$). This autocorrelation exhibits only one timescale that is consistent across networks with different connectivity structure. The global timescale is equal to the slowest interaction timescale in the autocorrelations of local activity related to the zero spatial frequency mode: $\tau_{\text{global}} = (1 - p_{\text{s}} - 8p_{\text{r}})^{-1}$ (Methods). However, this timescale has a very small relative contribution in local autocorrelations (scaled with the inverse number of neurons in the network) and is hard to observe empirically as it requires data with excessively long trial duration.

The relation between the spatial network structure and the timescales of local dynamics is a general phenomenon that is observed in the networks with linear (Fig. 6b) and non-linear (Supplementary Fig. 12, Methods) dynamics and different sizes (Supplementary Fig. 13). Moreover, this relation is independent of the number of connections per unit as long as they are uniformly distributed within the connectivity radius and their strength is normalized to the same total strength Supplementary Fig. 14, details in [10]).

#### 1.6 Local timescales in spatial network models with different interaction types

Interactions between units in the network model with transition rates described by Eq.(20) can be defined in different ways. To explain the presence of multiple timescales in the autocorrelation of neural activity, we model local, short-range connections between cortical minicolumns ($< 1$ mm) with strong, mean-driven interactions. For this mean-driven interaction type, the state of each unit $S_{x,y} \in \{0, 1\}$ is directly affected by the state of neighboring units $S_{x\pm1,y\pm1}$

$$S_{x,y} \leftarrow \beta[S_{x\pm1,y\pm1}], \tag{54}$$

where $\beta$ is the strength of interactions. On the other hand, it is also possible to use diffusion-type interactions. For diffusion-type interactions, the state of each unit is updated based on the difference between its state and the states of neighboring units

$$S_{x,y} \leftarrow \beta[S_{x\pm1,y\pm1} - S_{x,y}]. \tag{55}$$

This interaction type was used to model the long-range connections ($> 1$ mm) between cortical columns [13].

These different types of interactions create different spatiotemporal dynamics that give rise to different interaction timescales. Using Eq.(28) for the time-delayed quadratic moment in the Fourier space, we can write the spatiotemporal correlations $CC(k)$ for the network with the mean-driven interactions as

$$\tau_0 \frac{d}{dt} CC(k) \approx -[1 - \frac{\beta_1}{\alpha_1 + \alpha_2}(n - k^2)CC(k) + \ldots]. \tag{56}$$

Here $t$ is the time lag, $k$ is the spatial frequency mode, $n$ is the number of connected neighbors per unit and $\tau_0$ is the self-excitation timescale. From this equation, we can compute the interaction timescales as

$$\tau_k = \frac{\tau_0}{1 - n\frac{\beta_1}{\alpha_1+\alpha_2}(1 - k^2/n)}. \tag{57}$$

We can see that the interaction timescales are larger than the self-excitation timescale, and the largest interaction timescale is

$$\tau_{k=0} = \frac{\tau_0}{1 - n\frac{\beta_1}{\alpha_1+\alpha_2}}. \tag{58}$$

Because of this hierarchical relation between the interaction timescales and the self-excitation timescale, the autocorrelation of a single unit in the model with mean-driven interactions exhibits multiple dominant timescales (similar to neural data autocorrelations).

The spatiotemporal correlations in the network with diffusion-type interactions are given by

$$\tau_0 \frac{d}{dt} CC(k) \approx -[1 + \frac{\beta_1}{\alpha_1 + \alpha_2} k^2 CC(k) + \ldots]. \tag{59}$$

In this case, the interaction timescales are computed as

$$\tau_k = \frac{\tau_0}{1 + \frac{\beta_1}{\alpha_1+\alpha_2}k^2}. \tag{60}$$

We can see that the interaction timescales in the model with diffusion-type interactions are smaller than the self-excitation timescale, such that the largest interaction timescale is equal to the self-excitation timescale

$$\tau_{k=0} = \tau_0. \tag{61}$$

Therefore, autocorrelation of a single unit in this model exhibits only one dominant timescale which is the self-excitation timescale.

#### 2 Supplementary tables

| Fixed effect | $\tau_{2,\text{att-in}}$ | $\tau_{2,\text{att-away}}$ | $\tau_{1,\text{att-in}}$ | $\tau_{1,\text{att-away}}$ |
|---|---|---|---|---|
| Fixed effect, slope ($\omega_1 \pm 95\%$ CIs) | $-0.16 \pm 0.066$ | $0.015 \pm 0.12$ | $0.0016 \pm 0.86$ | $0.53 \pm 0.94$ |
| Fixed effect, intercept ($\omega_0 \pm 95\%$ CIs) | $343.76 \pm 53.95$ | $301.98 \pm 65.05$ | $308.35 \pm 62.10$ | $299.17 \pm 62.76$ |
| P-value for $\omega_1$ | $9 \times 10^{-6}$ | 0.79 | 0.997 | 0.26 |
| Monkey G, intercept ($\Omega_{0,G} \pm 95\%$ CIs) | $-35.53 \pm 52.62$ | $-43.3 \pm 63.25$ | $-47 \pm 62.2$ | $-43.19 \pm 62.75$ |
| Monkey B, intercept ($\Omega_{0,B} \pm 95\%$ CIs) | $-25.22 \pm -52.41$ | $-28.83 \pm 63.28$ | $-24 \pm 62.2$ | $-28.42 \pm 62.75$ |
| Monkey R, intercept ($\Omega_{0,R} \pm 95\%$ CIs) | $60.75 \pm 54.06$ | $72.13 \pm 64.09$ | $71 \pm 63.81$ | $71.61 \pm 63.52$ |
| Number of observations | 58 | 32 | 58 | 32 |
| F statistics | 23.87 | 0.07 | $1.36 \times 10^{-5}$ | 1.30 |
| Degrees of freedom (DF1, DF2) | $(1, 56)$ | $(1, 30)$ | $(1, 56)$ | $(1, 30)$ |
| $R^2$, adjusted | 0.62 | 0.69 | 0.46 | 0.70 |
| Cohen's $f^2$, effect size | 1.64 | 2.21 | 0.84 | 2.35 |

**Supplementary Table 1.** Results of fitting a mixed-effects model with a single fixed effect parameter (fast or slow timescale) and three random effects (for three monkeys: G, B, R) to quantify the relation between monkeys' reaction times ($RT$) and MAP estimates of timescales. Each column indicates parameters for one model as $RT_{i,m} = \omega_0 + \omega_1 \tau_{j,\text{cond},i} + \Omega_{0,m} + \varepsilon_{i,m}$, where $j \in \{1, 2\}$ indicates the fast (1) or slow (2) timescale, cond indicates attention condition (attend-in or attend-away) and $m$ indicates monkey. The model in MATLAB 2021a notation: $RT \sim 1 + \tau_{j,\text{cond}} + (1|\text{Monkey})$.

| Attention condition | Attend-in | Attend-away |
|---|---|---|
| Coefficient for $\tau_2$ ($\omega_1 \pm 95\%$ CIs) | $-0.16 \pm 0.066$ | $-0.0058 \pm 0.12$ |
| Coefficient for $\tau_1$ ($\omega_2 \pm 95\%$ CIs) | $0.34 \pm 0.73$ | $0.54 \pm 1.00$ |
| Fixed effect, intercept ($\omega_0 \pm 95\%$ CIs) | $341.63 \pm 54.07$ | $299.90 \pm 64.60$ |
| P-value for $\omega_1$ | $6.02 \times 10^{-6}$ | $0.92$ |
| P-value for $\omega_2$ | $0.36$ | $0.27$ |
| Monkey G, intercept ($\Omega_{0,G} \pm 95\%$ CIs) $\pm 95\%$ CIs | $-36.94 \pm 52.65$ | $-43.15 \pm 62.73$ |
| Monkey B, intercept ($\Omega_{0,B} \pm 95\%$ CIs) | $-23.56 \pm 52.48$ | $-28.32 \pm 62.76$ |
| Monkey R, intercept ($\Omega_{0,R} \pm 95\%$ CIs) | $60.50 \pm 53.98$ | $71.46 \pm 63.56$ |
| Number of observations | 58 | 32 |
| F statistics | 12.54 | 0.65 |
| Degrees of freedom (DF1, DF2) | $(2, 55)$ | $(2, 29)$ |
| $R^2$, adjusted | 0.62 | 0.69 |
| Cohen's $f^2$, effect size | 1.64 | 2.24 |

**Supplementary Table 2.** Results of fitting a mixed-effects model with two fixed effect parameters (fast and slow timescales) and three random effects (for three monkeys: G, B, R) to quantify the relation between monkey's reaction times ($RT$) and MAP estimates of timescales. Each column indicates parameters for one model as $RT_{i,m} = \omega_0 + \omega_1 \tau_{2,\text{cond},i} + \omega_2 \tau_{1,\text{cond},i} + \Omega_{0,m} + \varepsilon_{i,m}$, where cond indicates attention condition (attend-in or attend-away) and $m$ indicates monkey. The model in MATLAB 2021a notation: $RT \sim 1 + \tau_{2,\text{cond}} + \tau_{1,\text{cond}} + (1|\text{Monkey})$.

### 3 Supplementary figures

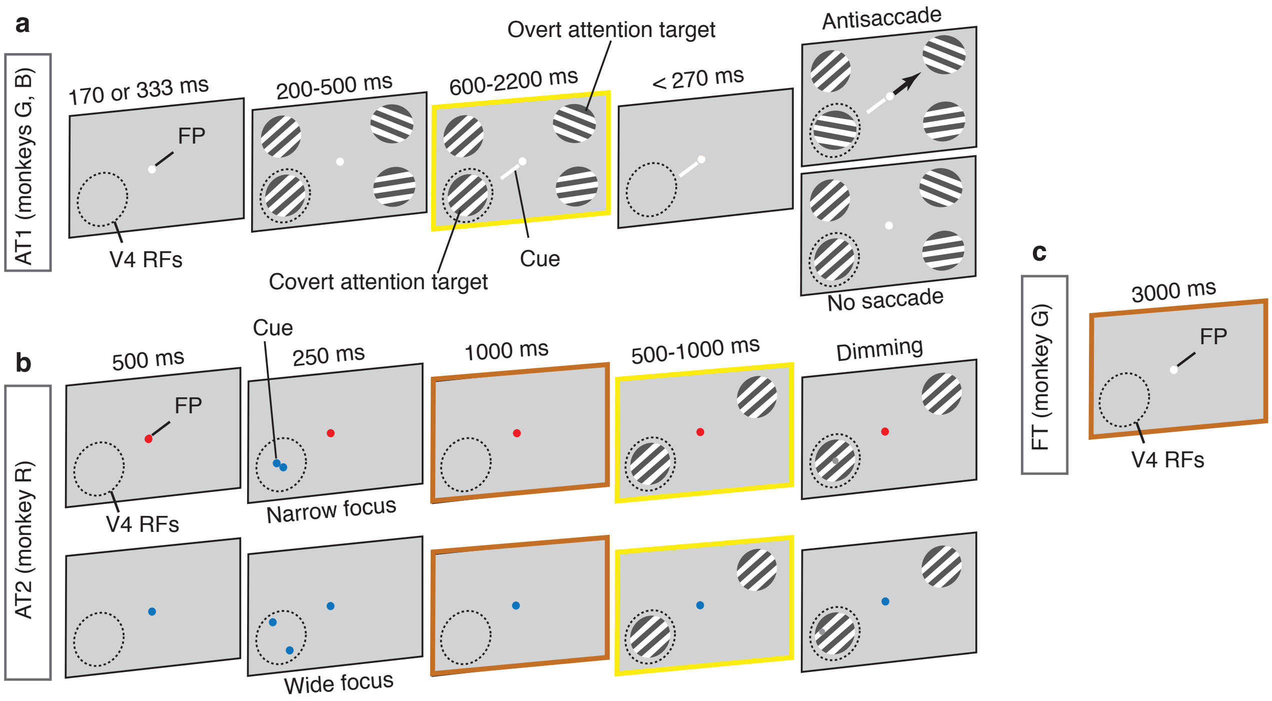

**Supplementary Fig. 1. Behavioral tasks. (a)** Attention task 1 (AT1). Monkeys started a trial by fixating a central dot (FP). Following a brief delay (333 ms and 170 ms in monkeys G and B, respectively), four peripheral oriented-grating stimuli were presented. After a variable delay, stimuli disappeared then reappeared, either with or without one of the four stimuli rotating. Monkeys reported the rotation by making a saccade to the stimulus diametrically opposite to the change (Antisaccade; arrow shows the saccade direction) or maintained fixation if no rotation happened (No saccade). A small, central cue (white line) indicated which stimulus was likely to change. The dashed circle, denoting the V4 receptive fields locations (V4 RFs), and saccade direction arrow were not visible to the monkeys. **(b)** Attention task 2 (AT2). The monkey initiated a trial by holding a bar and visually fixating the fixation point. The color of the fixation point indicated the level of spatial certainty (red: Narrow focus, blue: Wide focus). Shortly after, a cue appeared indicating the area (but also the focus) of the visual field to attend to. One second after the disappearance of the cue, two gratings appeared. One in the center of the RF and one diametrically opposed along the fixation point. The monkey was rewarded for detecting a small luminance change (Dimming, gray point) that occurred either in the center of the grating (narrow focus) or in one of peripheral positions (wide focus) by releasing the bar. **(c)** Fixation task (FT). The monkey was rewarded for fixating a central dot on a blank screen for 3 s on each trial. In all tasks, epochs marked with brown frames were used for spontaneous nerual activity and epochs marked with yellow frames were used for stimulus-driven activity. Graphical elements are not precisely to scale.

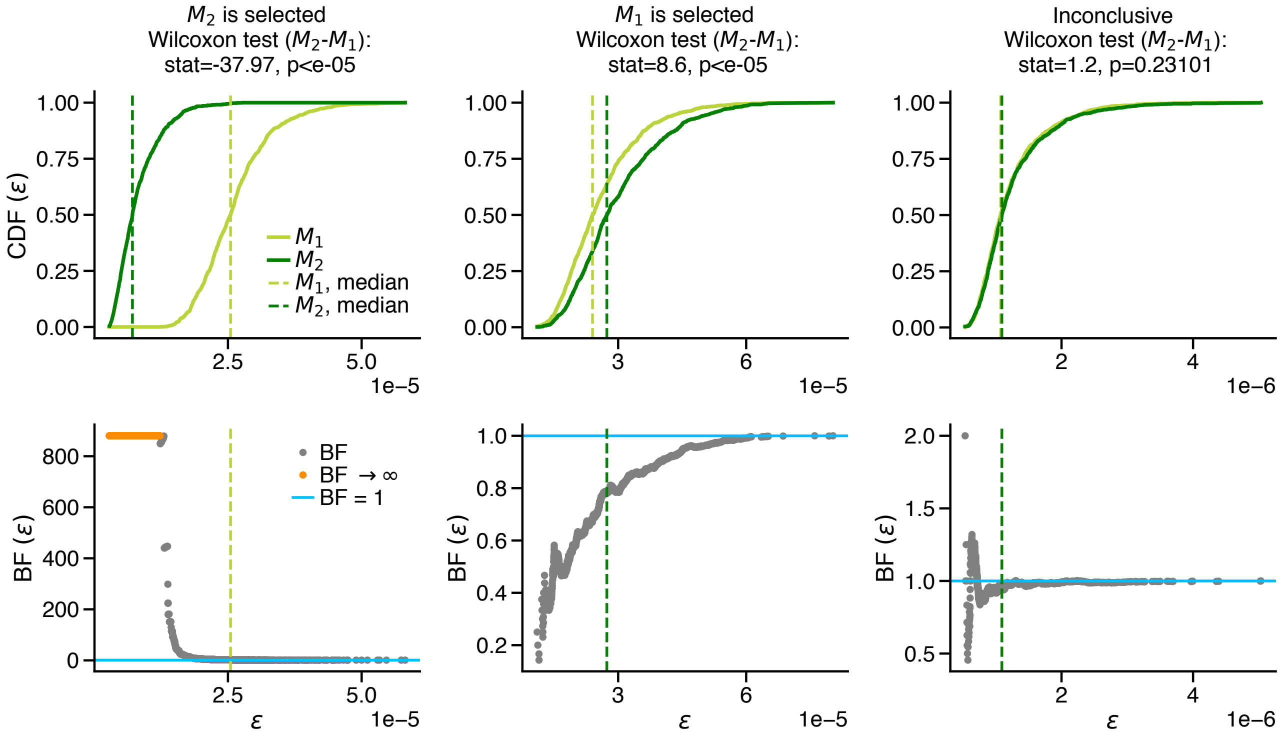

**Supplementary Fig. 2. Examples of different outcomes of the Bayesian model comparison.** We use the CDFs of distance distributions for the two-timescale ($M_2$) and one-timescale ($M_1$) models (upper row) to approximate the Bayes factors $\text{BF}(\varepsilon) = \text{CDF}_{M_2}(\varepsilon)/\text{CDF}_{M_1}(\varepsilon)$ for different values of error threshold ($\varepsilon$, lower row). If the distance distributions of the two models were significantly different (Wilcoxon ranksum test) and $\text{BF}(\varepsilon) > 1$ for all $\varepsilon < \max_{M_1,M_2}[\text{median}(\varepsilon)]$ (dashed vertical lines), then $M_2$ was selected (left panels), and if $\text{BF}(\varepsilon) < 1$, then $M_1$ was selected (middle panels). Otherwise the model comparison outcome was categorized as inconclusive (right panels). Orange dots in the lower right panel indicate the error thresholds where the Bayes factor diverged due to $\text{CDF}_{M_1}(\varepsilon) = 0$.

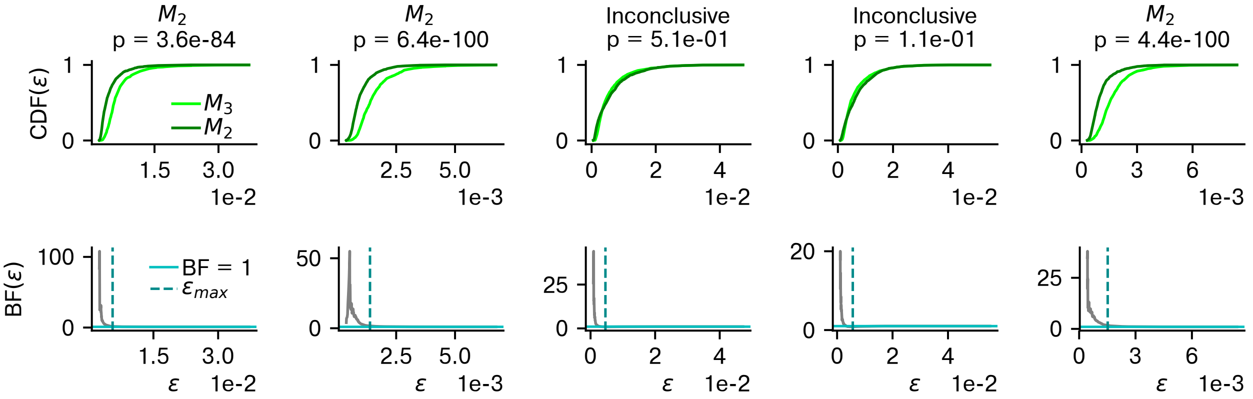

**Supplementary Fig. 3. Model comparison between two- and three-timescale models.** Three-timescale model did not provide a better description of V4 autocorrelations than the two-timescale model. The Bayesian model comparison between two- ($M_2$) and three-timescale ($M_3$) models selects $M_2$ model ($\mathrm{BF}(\varepsilon) = \mathrm{CDF}_{M_2}(\varepsilon)/\mathrm{CDF}_{M_3}(\varepsilon) > 1$, $\forall\varepsilon < \varepsilon_{\mathrm{max}}$, $\varepsilon_{\mathrm{max}} = \max_{M_2,M_3}[\mathrm{median}(\varepsilon)]$) in $3$ out of $5$ sessions in the fixation task, and the outcome for the $2$ remaining sessions is inconclusive. Upper row: CDFs of distance distributions for the two models. Lower row: Corresponding Bayes factors. Panel titles indicate the outcome of the model comparison with p-values estimated from a two-sided Wilcoxon rank-sum test. Data are from the fixation task (FT) in monkey G which had the longest trial duration ($3$ s).

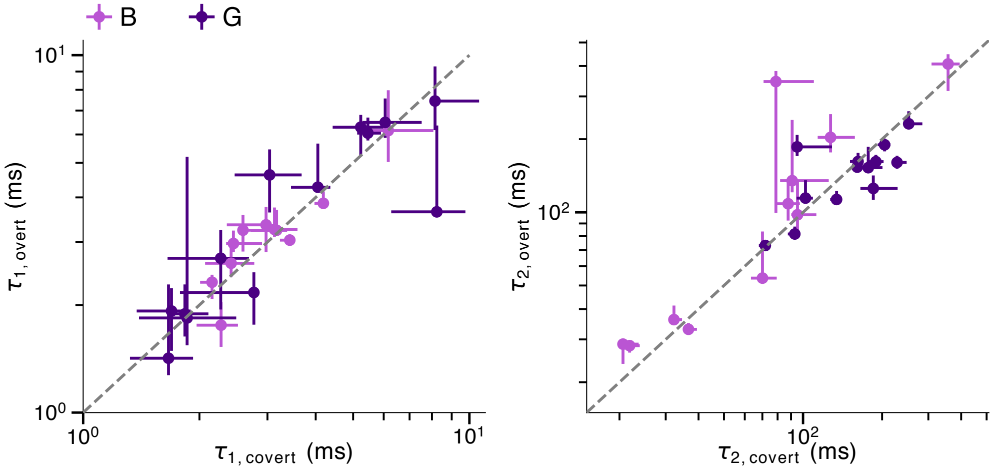

**Supplementary Fig. 4. Comparing timescales between covert and overt attention.** There was no significant difference in fast (left) and slow (right) timescales between the covert and overt attention conditions ($p > 0.05$ for both $\tau_1$ and $\tau_2$, two-sided Wilcoxon signed-rank test). Points - MAP estimates for individual sessions, error bars - the first and third quartiles of the marginal posterior distribution, dashed line - the unity line. If the MAP estimate was smaller than the first or larger than the third quartile, the error bar was discarded. Color of the dots indicates different monkeys.

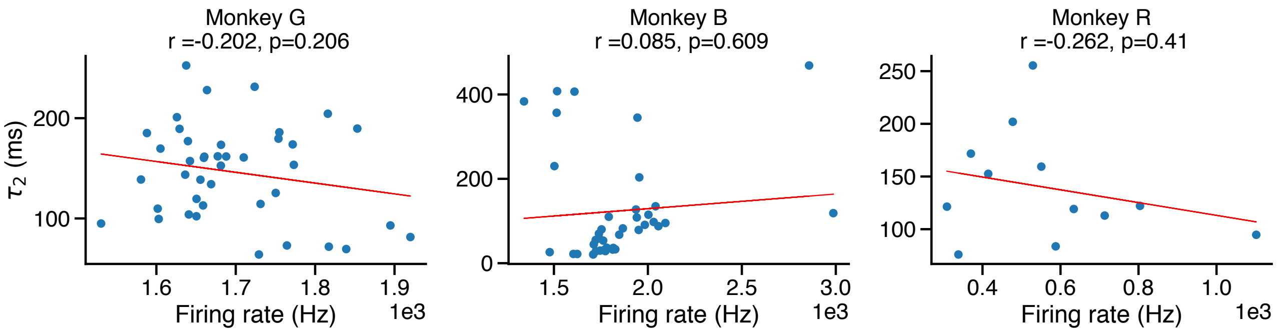

**Supplementary Fig. 5. The slow timescale ($\tau_2$) does not depend on the mean firing rate across sessions and attention conditions.** The MAP estimates of $\tau_2$ from aABC fits versus the overall mean firing rate of the population activity for individual sessions and attention conditions (red line - linear regression line). $r$ is the Pearson correlation coefficient and $p$ is the corresponding p-value.

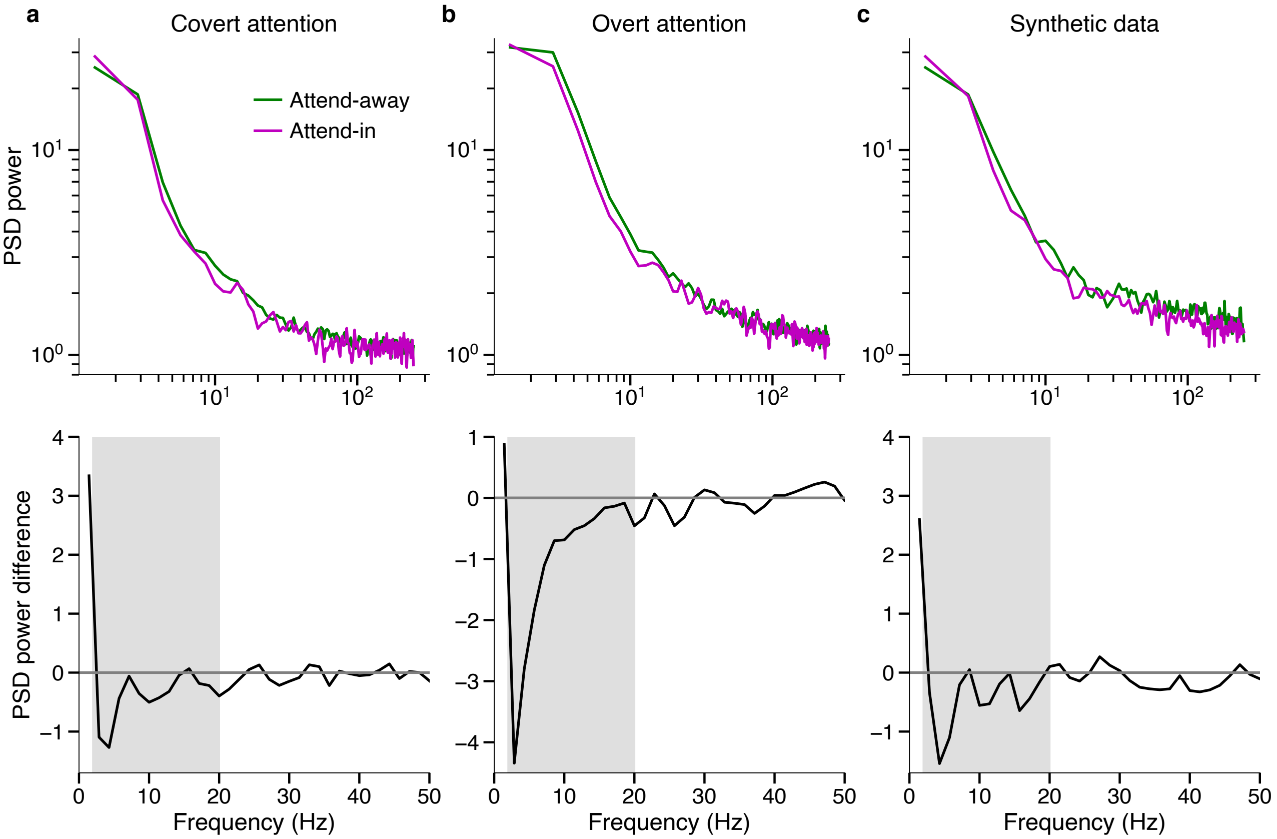

**Supplementary Fig. 6. Reduction in the power of low-frequency fluctuations during attention. (a)** The power spectral densities (PSDs, upper row) of spiking activity for covert attention and attend-away condition (same data as in Fig. 3a) and their difference (attend-in $-$ attend-away, lower row). The power is reduced during attention within the frequency range $\sim 2 - 20$ Hz (gray shading) as reported in previous studies[14, 1, 2, 4]. **(b)** Same as a for overt attention (same data as in Fig. 3d). **(c)** Synthetic data generated from a two-timescale generative model with MAP parameters estimated from data in a exhibit a similar PSD shape as V4 data. The increase in the slow timescale during attention predicts an increase in the spectral power at very low frequencies ($< 2$ Hz), which is not detectable with short observation windows ($< 500$ ms) used in previous studies.

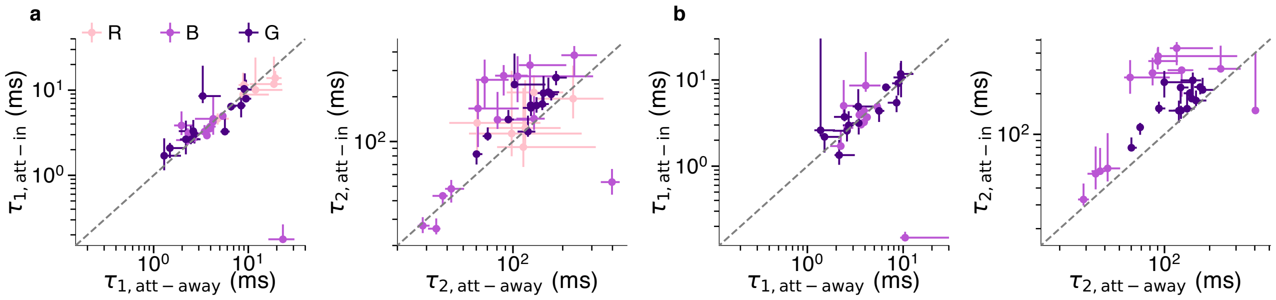

**Supplementary Fig. 7. Timescales estimated by fitting the power spectral density of spiking activity over the entire range of frequencies during attention tasks. (a)** Slow timescales ($\tau_2$, right) increase during covert attention (mean $\tau_{2,\text{att-in}} = 166.39$ ms, mean $\tau_{2,\text{att-away}} = 120.73$ ms, $p = 2 \times 10^{-4}$, $N = 32$, two-sided Wilcoxon signed-rank test). Points - MAP estimates for individual sessions, error bars - the first and third quartiles of the marginal posterior distribution, dashed line - the unity line. If the MAP estimate was smaller than the first or larger than the third quartile, the error bar was discarded. Color of the dots indicates different monkeys. **(b)** Same as a for the overt attention (mean $\tau_{2,\text{att-in}} = 199.29$ ms, mean $\tau_{2,\text{att-away}} = 121.90$ ms, $p = 10^{-4}$, $N = 26$, two-sided Wilcoxon signed-rank test). The results agree with timescales estimated in time-domain (Fig. 3).

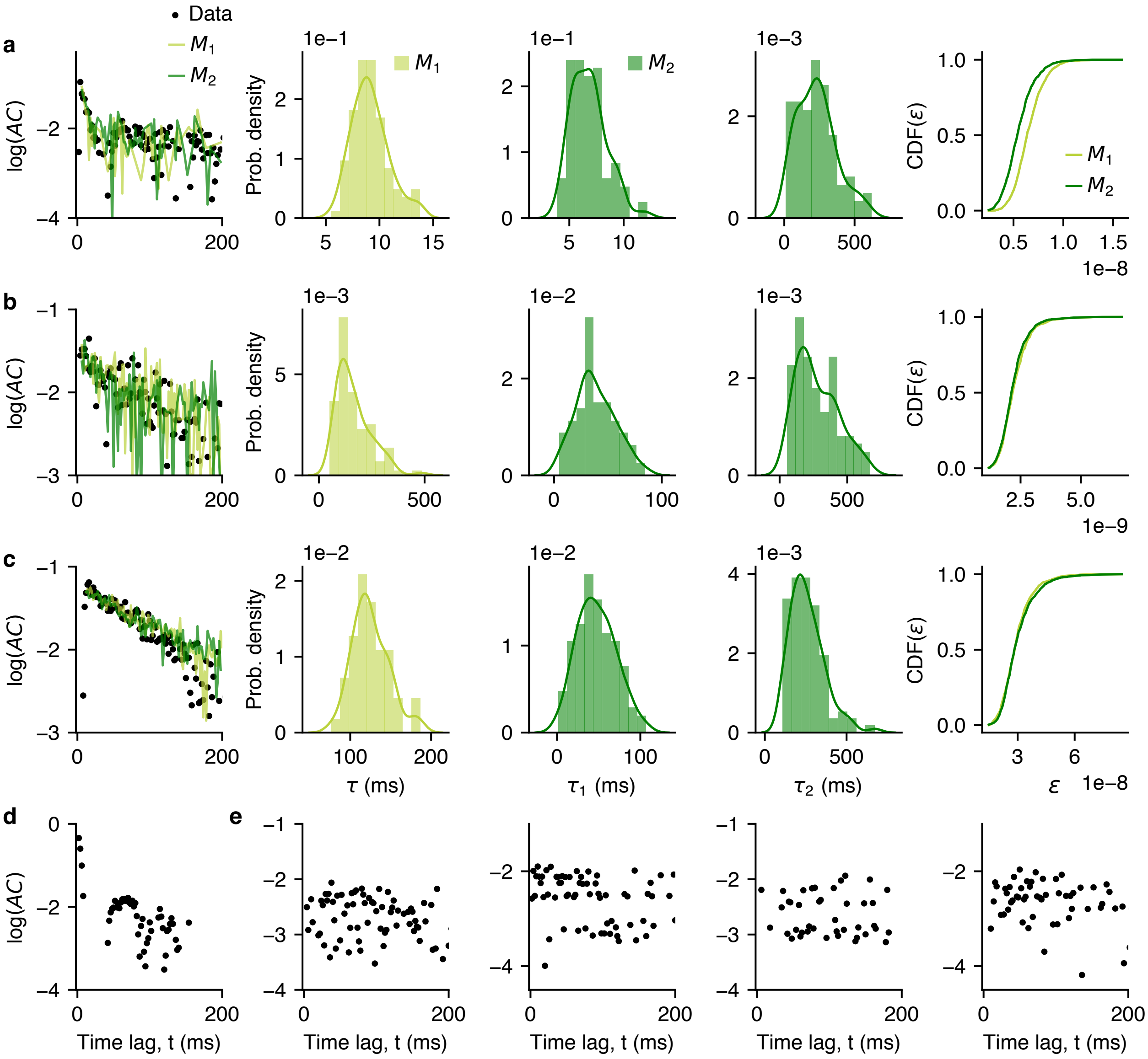

**Supplementary Fig. 8. Shape and timescales of single-unit autocorrelations. (a)** Autocorrelation of an example single-unit activity that is better fitted with the two-timescale model ($M_2$) than the one-timescale model ($M_1$). Left: Single-unit autocorrelation overlaid by the autocorrelations of synthetic data from $M_2$ and $M_1$ with MAP parameters and the same statistics (spike rate and variability, the number and duration of trials) as in the experimental data. Data autocorrelations are plotted from the first time-lag ($t = 2$ ms). Middle: Marginal posterior distributions of the timescales estimated by fitting $M_1$ ($\tau$) or $M_2$ ($\tau_1$ and $\tau_2$). Right: Cumulative distribution of errors $\text{CDF}_{M_i}(\varepsilon)$ between the autocorrelations of the single-unit data and synthetic data generated with parameters sampled from the $M_1$ or $M_2$ posteriors. $M_2$ is a better fit since it produces smaller errors (i.e. Bayes factor $= \text{CDF}_{M_2}(\varepsilon)/\text{CDF}_{M_1}(\varepsilon) > 1$, Methods). **(b, c)** Same as a for example single units with an inconclusive outcome of the model selection, i.e. there is no significant difference between the distributions of errors for the two models. **(d)** Example single-unit autocorrelation dominated by an oscillation. **(e)** Examples of single-unit autocorrelations dominated by statistical bias without any clear temporal structure due to low firing rate.

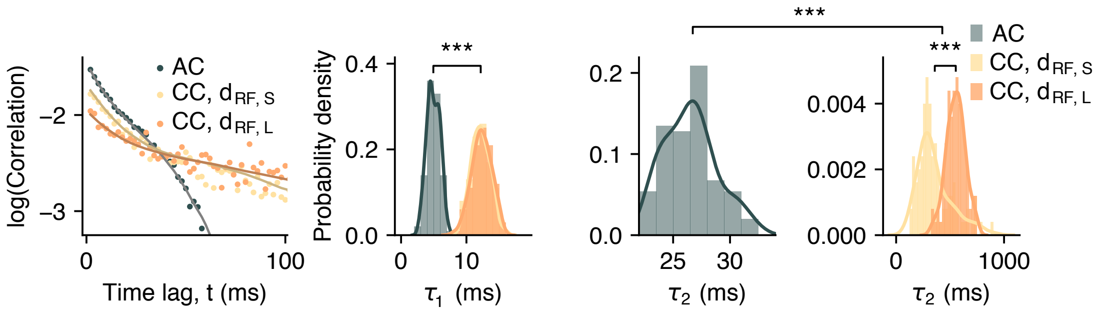

**Supplementary Fig. 9. Timescales of auto- and cross-correlations of V4 neurons.** Auto- and cross-correlations of V4 neurons recorded across different electrode channels (left, AT2) and corresponding posterior distributions of timescales (right) (data from monkey R, similar to Fig. 5j,k in the main text for monkey G). The shape of auto- and cross-correlations is captured by the autocorrelation of synthetic data from the two-timescale OU process with MAP parameters (fitted lines, Methods). The strengths of cross-correlations is smaller than the auto-correlation and decreases with RF-center distance (left, $d_{\text{RF,L}} > d_{\text{RF,S}}$). Cross-correlations have slower timescales than the autocorrelation. The slow timescale in cross-correlations increases with increasing RF-center distance, but there is no significant difference in the fast timescale (right) (different from Fig. 5k in the main text for monkey G, where there was a significant increase also in the fast timescale). Statistics: two-sided Wilcoxon rank-sum test, *** indicates $p < 10^{-3}$. Number of samples in each posterior $N = 100$. Correlations are plotted from the first time-lag ($t = 2$ ms).

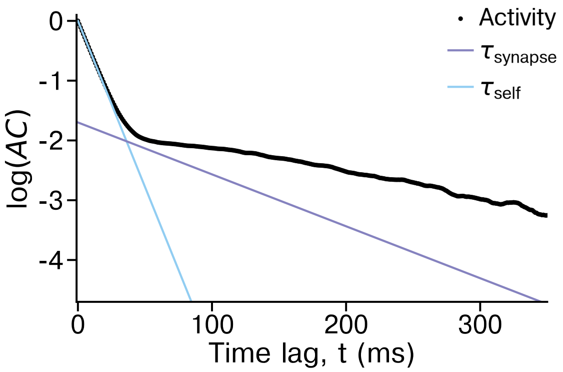

**Supplementary Fig. 10. Timescales in spatial network with synaptic filtering.** Autocorrelation of individual units' activity in a network model with local spatial connectivity and synaptic filtering exhibits at least two distinct timescales. The slow timescale is larger than the synaptic timescale ($\tau_{\text{synapse}}$) due to local recurrent interactions.

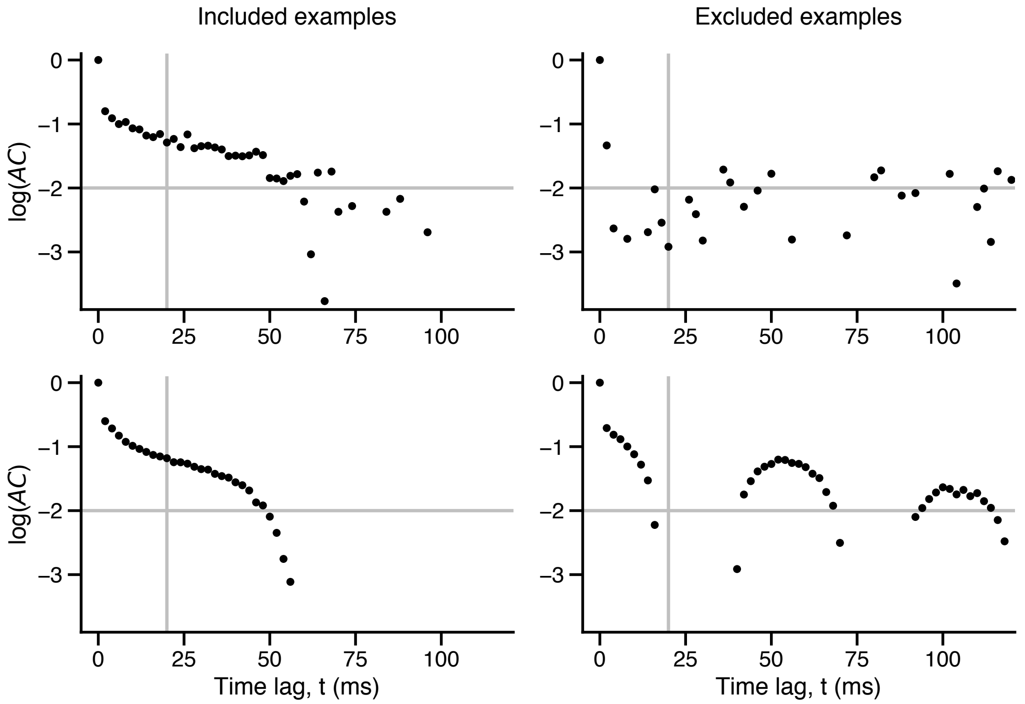

**Supplementary Fig. 11. Examples of included (left) and excluded autocorrelations (right) for timescales estimations.** We excluded autocorrelations dominated by noise (upper right) or strong oscillations (lower right) that could not be well described with a mixture of exponential decay functions. The autocorrelation was excluded if it fell below $0.01$ ($\log(AC)$ fell below $-2$) in lags smaller or equal to $20$ ms (gray lines). The drop between the zero time-lag and the first time-lag ($t = 2$ ms) in all examples reflects the difference between the total variance of spike counts and the variance of instantaneous rate according to the law of total variance.

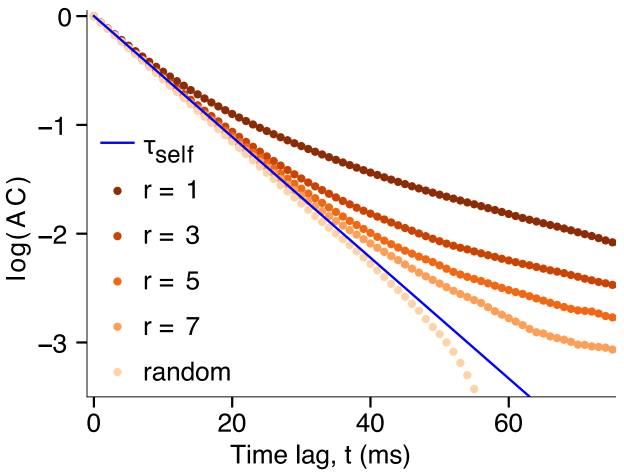

**Supplementary Fig. 12. Local timescales in network models with spatial connectivity and non-linear interactions reflect the underlying network structure.** In the network with non-linear interactions (Eq. 13 in the main text), local autocorrelations become dominated by the self-excitation timescale ($\tau_{\text{self}}$) when increasing the connectivity radius while keeping the connection strengths constant ($p_{\text{s}} = 0.88, 8p_{\text{r}} = 0.12$). This result is similar to the network with linear interactions (cf. Fig. 6, Eq. 12 in the main text) .

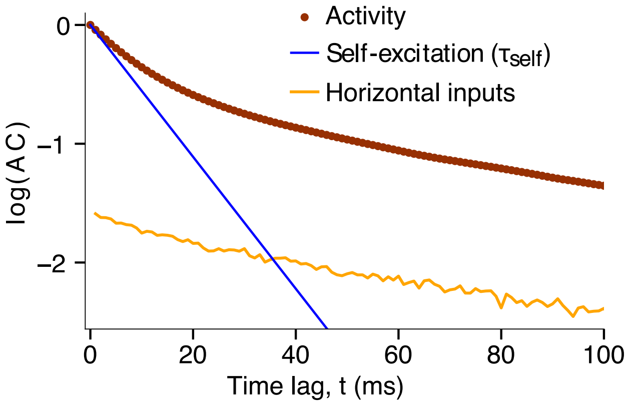

**Supplementary Fig. 13. Multiple timescales in autocorrelations of spatially connected networks of different size.** Autocorrelation (AC, brown) of individual units' activity in a network with $100$ units ($10 \times 10$ lattice) exhibits multiple timescales that are similar to the timescales in a network with $10,000$ units ($100 \times 100$, shown in Fig. 5f in the main text). The fast timescale ($\tau_{\mathrm{self}}$) in initial time-lags can be estimated by the autocorrelation of a 2-state Markov process driven only by $p_{\mathrm{s}}$ and $p_{\mathrm{ext}}$ (blue line). Slower timescales in larger time-lags can be captured by the autocorrelation of horizontal inputs to each unit (orange line).

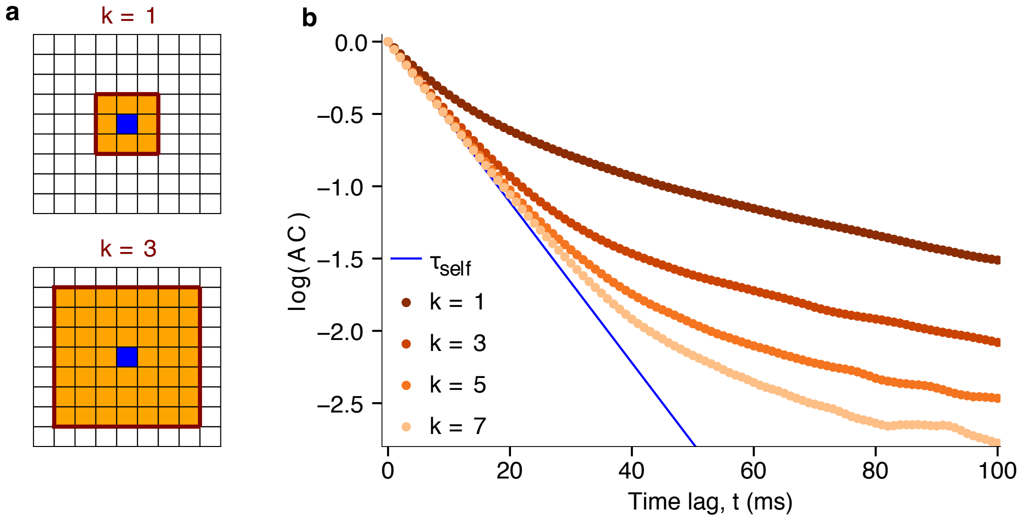

**Supplementary Fig. 14. Local timescales in a network model with dense local connectivity. (a)** Schematics of dense local connectivity with small ($k = 1$) and larger ($k = 3$) connectivity radius. Each unit (blue) is connected to all units (orange) within the connectivity radius $k$ (brown square). For different connectivity radius, we keep the total connection strength ($np_{\mathrm{r}}$, $n$ is the number of connections per unit) constant. **(b)** For different connectivity radii, the shape of autocorrelations of individual units ($AC$) reflects the underlying spatial connectivity (cf. Fig. 6 in the main text). Local autocorrelations become dominated by the self-excitation timescale ($\tau_{\mathrm{self}}$) when increasing the connectivity radius while keeping the connection strengths constant ($p_{\mathrm{s}} = 0.88, np_{\mathrm{r}} = 0.11$).
